# Characterization of *ORF19.7608* (*PPP1*), a Biofilm-induced Gene of *Candida albicans*

**DOI:** 10.1101/2025.06.28.660404

**Authors:** Nmerichukwu C. Iwuchukwu, Anna Carolina Borges Pereira da Costa, Chris Law, Min-Ju Kim, Aaron P. Mitchell, Malcolm Whiteway

**Affiliations:** Department of Biology, Concordia University, Montreal, Quebec, Canada; Centre for Microscopy and Cellular Imaging, Concordia University, Montreal, Quebec, Canada; Department of Microbiology, University of Georgia, Athens, Georgia, USA

## Abstract

The opportunistic human pathogen *Candida albicans* is an important cause of nosocomial infections, in large part because of its propensity to form biofilms on indwelling medical devices such as catheters. The formation of these biofilms is controlled by a complex transcriptional network and involves over a thousand genes, many of which are uncharacterized. We have investigated three genes (*ORF19.4654*, *ORF19.7608,* and *PBR1*), found only in *C. albicans* and closely related species, that are highly induced under biofilm conditions and encode small proteins with N-terminal signal sequences. Through the construction of fluorescent protein fusions, we have examined the location of the encoded proteins in both planktonic and biofilm cells. Orf19.4654-Scarlet and Pbr1-Scarlet were localized to the vacuole under both conditions. In contrast, the Orf19.7608-GFP fusion generated a punctate pattern only under biofilm conditions and was designated Ppp1 (Punctate Pattern Protein 1). The Ppp1-GFP puncta were similar in location, stability, and size to those formed by the eisosome subunit Sur7, but co-localization studies suggest that Ppp1 and Sur7 define separate elements. The *PPP1* mutation does not cause a distinct phenotype under various stress conditions or in the presence of antifungals and does not impact biofilm formation and biomass. These data suggest that while the expression and cellular localization of Ppp1 appear controlled by conditions generating biofilms, and define a unique subcellular localization pattern, Ppp1 protein function is not essential for biofilm formation.

**IMPORTANCE:** Biofilm formation is a virulence factor of medical importance in *C. albicans*. Identifying the biological function and cellular localization of biofilm-related proteins can help in their characterization and understanding of the biofilm network. Microscopy was employed to identify the subcellular localization of putative biofilm proteins, aiming to provide insight into their possible functions. Prb1 and Orf19.4654 localized to the vacuole, while Orf19.7608 (Ppp1) formed puncta. Phenotypic assays to investigate the uniqueness of *PPP1* revealed that it does not play a major role in the stress response pathways or antifungal activity. Overall, our study provides insight into the localization of the products of Candida-specific biofilm genes.

## INTRODUCTION

*Candida albicans* is a part of the normal human microbial flora and is typically found in the mouth, skin, gastrointestinal tract, and vagina (Odds 1988, Kumamoto 2011). However, when the body’s pH is altered, the microbiota composition is changed, or the immune system is compromised, this commensal organism can become a pathogen (Correia et al., 2015). The changes in the interaction with its host, from commensal to pathogenic, can be attributed to various environmental factors, cellular processes, genetic responses, and induction pathways (Pemmaraju et al., 2016; Kumar and Kumar, 2023). Virulence factors include hydrolytic enzymes, genome plasticity, adherence and invasion enzymes, metabolic plasticity, heat shock proteins, and antifungal resistance. In particular, biofilm formation has a significant influence on *C. albicans* as a nosocomial pathogen (Mba and Nweze 2020).

A biofilm is a community of microorganisms attached to a surface and enclosed within an extracellular matrix (Lewis, 2001; Donlan, 2001). Biofilm-forming microorganisms form colonies of cells that can resist the host’s immune response as well as antimicrobial drug treatments. The adherence of biofilms to surfaces, such as catheters and prostheses, makes them a significant contributor to hospital-acquired infections (Wisplinghoff et al., 2003). Biofilm cells are usually resistant to antifungals and host immunity, so they pose a huge problem in healthcare (Mukherjee and Chandra 2004, Kaur and Nobile 2023).

*C. albicans* biofilm formation is regulated by environmental factors like osmotic stress, oxidative stress, and heat shock, as well as genetic factors like secondary messengers, transcription factors, and regulatory RNAs (Pemmaraju et al., 2016; Kumar and Kumar, 2023). Osmotic and oxidative stress play an important role in fungal virulence, with osmotic stress increasing the tonicity of the cell, thereby causing water loss and cell size reduction, and oxidative stress causing the release of reactive oxygen species (ROS) (Pemmaraju et al., 2016; Kühn and Klipp, 2011). Other factors, such as drug efflux pumps, quorum sensing, host immunity, and extracellular matrix (ECM) also influence biofilm formation (Kaur and Nobile 2023).

A comprehensive study has investigated the transcriptional network regulating *C. albicans* biofilms (Nobile et al., 2012). More than 1000 genes were identified, many of which have already been extensively studied. However, some highly up-regulated genes are poorly characterized; these include the genes *ORF19.7608*, *ORF19. 4654* and *PBR1,* which encode N-terminal signal peptide-containing proteins found only in *C. albicans* and its closest relatives, *C. dubliniensis* and *C. tropicalis*.

Signal peptides are short, typically hydrophobic amino acid sequences in nascent proteins associated with the secretory pathway, including proteins that are localized in the secretory organelles like the plasma membrane (PM), endoplasmic reticulum (ER), vacuole and Golgi, proteins that are secreted, and proteins localized in other subcellular compartments (Wild et al., 2004). They are N-terminal recognition sequences that help in the transport and secretion of proteins.

The plasma membrane (PM) is an integral cell organelle that forms a protective bilayer, regulating cellular processes to maintain cell homeostasis. It interacts with both the environment and the cell’s cytoplasm (containing other organelles), leading to a role in most cell functions from environmental sensing to nutrient uptake, cellular morphogenesis, secretion, endocytosis, and cell wall biogenesis (Alvarez et al., 2008; Douglas et al., 2011). Some genes involved in biofilm formation, such as adhesins, have a GPI anchor attaching them to the plasma membrane (Desai & Mitchell, 2015). Due to its important roles, it is the major target of antifungals, with two of the four classes of antifungals designed to inhibit its synthesis or function (Douglas & Konopka, 2016). The PM contains three distinct subdomains with similar protein distribution, namely the Membrane Compartment occupied by Can1 (MCC) or eisosome, the Membrane Compartment occupied by Pma1 (MCP), and the TORC2 complex /Membrane Compartment containing TORC2 (MCT) (Douglas et al., 2011; Malinsky et al., 2010).

Fluorescence microscopy allows visual examination of the patterns of subcellular distribution of biomolecules labelled with fluorescent probes such as green fluorescent protein (GFP) (Tsien, 1998). Labelling of subcellular compartments with complementary fluorescent probes allows the analysis of the co-occurrence of such probes, which can be used to determine the likelihood of a given protein being found in a given subcellular domain (Dunn et al., 2011). This co-occurrence can be quantified using Pearson’s Correlation Coefficient (PCC) or Manders’ Colocalization Coefficient (MCC) (Dunn et al., 2011). Time-lapse imaging allows the analysis of protein motility within cells, and when multiple proteins are imaged at the same time, this can give more information about whether proteins are localised to the same subcellular domain (Naseem et al., 2024).

This study examines three genes, highly upregulated in biofilm formation, encoding small N-signal-peptide-encoding Candida-specific proteins (Nobile et al., 2012). Fluorescence microscopy identified a specific punctate pattern under biofilm conditions for Orf19.7608, leading us to name it Punctate Pattern Protein 1 (Ppp1). *ORF19.6274* (*PBR1*) and *ORF19.4654* encoded proteins that showed vacuolar localization. Due to its interesting protein distribution pattern, phenotypic assays were employed to highlight the role of Ppp1 in cellular responses and impact on biofilm structure and mass. Deletion of Ppp1 resulted in no significant changes in biofilm formation, demonstrating that Ppp1 is not a major player in cellular stress responses and Candida biofilm structure and biomass.

## RESULTS

### Bioinformatics analysis of biofilm genes

A study by Nobile et al. 2012 into the transcriptional network of biofilm regulation in *C. albicans* identified more than 1000 genes upregulated in biofilm formation. Many of the highly upregulated genes have been extensively studied. We expanded the investigation into other genes, namely *ORF19.7608, ORF19.6274 (PBR1*), and *ORF19.4654.* These genes were selected because although they are highly upregulated in biofilm formation (8.9 log2fold for *ORF19.7608*, 8.8 log2fold for *PBR1,* and 7.2 log2fold for *ORF19.4654*), they have not been extensively studied (Nobile et al., 2012). Protein database searches with UniProt to provide insight into possible structural and functional annotations identified signal peptides in all three proteins, at positions 1-19 for *ORF19.7608* and *ORF19.4654,* and 1-21 for *PBR1* (See Figure 1). Identification of orthologs using NCBI blast against the RefSeq database identified orthologs only in *C. dubliniensis* for *PPP1* and *ORF19.4654,* and both *C. dubliniensis* and *C. tropicalis* for *PBR1*.

**Figure 1:**
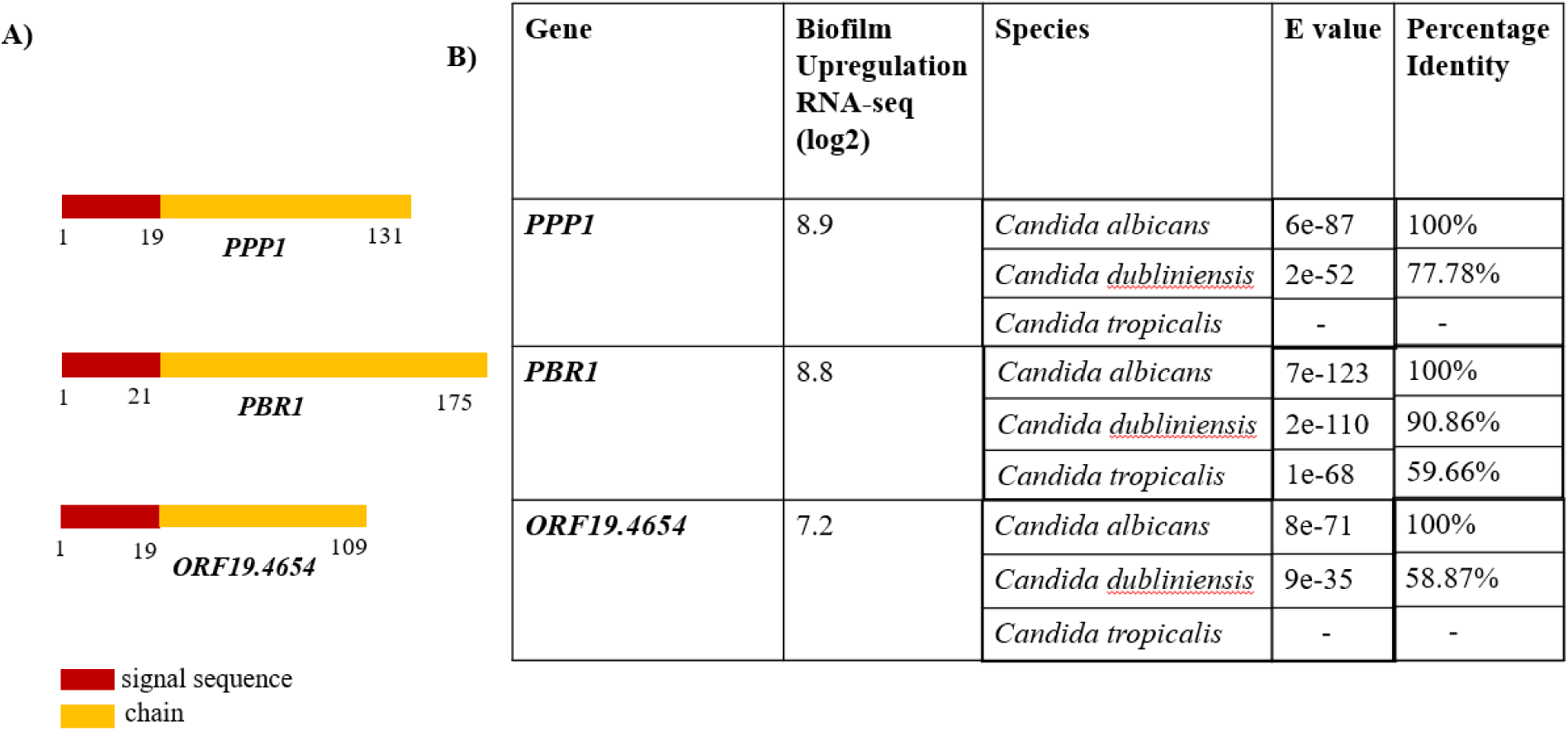
Bioinformatic analysis of three small biofilm genes found in only *Candida spp*. (A) Rectangles depicting the peptide composition of proteins screened from the biofilm library by Nobile et al. 2012. Red shows the N-signal peptide, and yellow shows the chain as predicted by UniProt (The UniProt Consortium 2023). (B) Table showing induction of biofilm genes and protein blast of amino acid sequences against the RefSeq database of proteins on NCBI.

### Fluorescence microscopy identifies a pattern for Orf19.7608 under biofilm conditions

Examining subcellular localization through fluorescence microscopy can help gain insight into a protein’s function as it links potential physiological processes to the protein through organellar compartmentalization (Combs & Shroff, 2017). We employed fluorescence microscopy to identify the cellular compartments where Orf19.7608, Pbr1, and Orf19.4654 are localized by making fusion constructs of the proteins, namely Orf19.7608-GFP, Pbr1-Scarlet, and Orf19.4654-Scarlet. Cells were grown under planktonic conditions for 16 hours and biofilm conditions for 48 hours to validate RNA seq data from Nobile et al. 2012 and check if growth conditions affect protein distribution. Under both planktonic and biofilm conditions, Pbr1 and Orf19.4654 have similar localization with protein distribution primarily in a large subcellular compartment (Figure 2). Orf19.7608, however, has a very different protein distribution, and this is dependent on growth conditions. In planktonic cells, the protein encoded by Orf19.7608 disperses in the cell, while evenly distributed cell periphery puncta are formed in yeast cells under biofilm conditions (Figure 2). Hyphal cells in biofilms do not show the same protein distribution as the yeast biofilm for any of the 3 proteins studied. The punctate distribution pattern formed the basis of the designation Punctate Pattern Protein 1 (Ppp1) for Orf19.7608.

**Figure 2:**
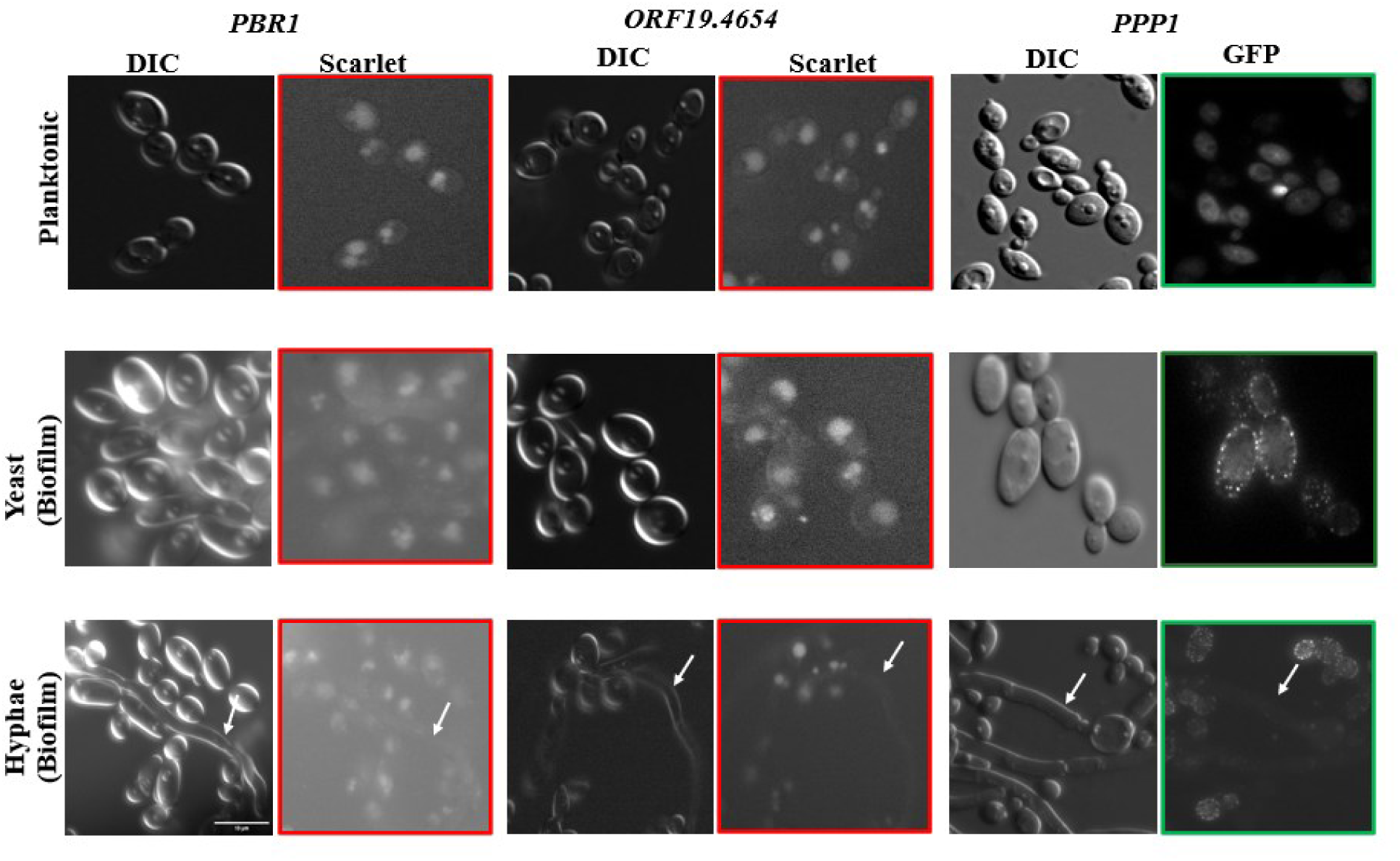
Fluorescence microscopy identifies a punctate pattern for Ppp1 in yeast biofilm cells. *PPP1-GFP*, *PBR1-Scarlet*, and *ORF19.4654-Scarlet* are grown in planktonic (YPD at 220rpm/30^0^C/16hrs) and biofilm conditions (Spider at 75rpm/37^0^C/48hrs) conditions. Under these conditions, Pbr1 and Orf19.4654 have the same protein distribution. However, Ppp1 forms puncta under yeast biofilm conditions. Arrows highlight hyphae cells. The scale bar represents 10µm.

### Pbr1::Scarlet and Orf19.4654::Scarlet localize to the vacuole

We further investigated the protein distribution of Pbr1 and Orf19.4654 to establish their specific sub-cellular localizations. Most of the signal for each protein was concentrated in a single large subcellular compartment, suggesting these proteins may be localised to the nucleus or the vacuole. This protein distribution corresponded with CMAC vacuolar staining, and not with staining of Hoechst 33342 for the nucleus (Figure 3), indicating that the Pbr1 and Orf19.4654 Scarlet fusion proteins localize in the vacuole.

**Figure 3:**
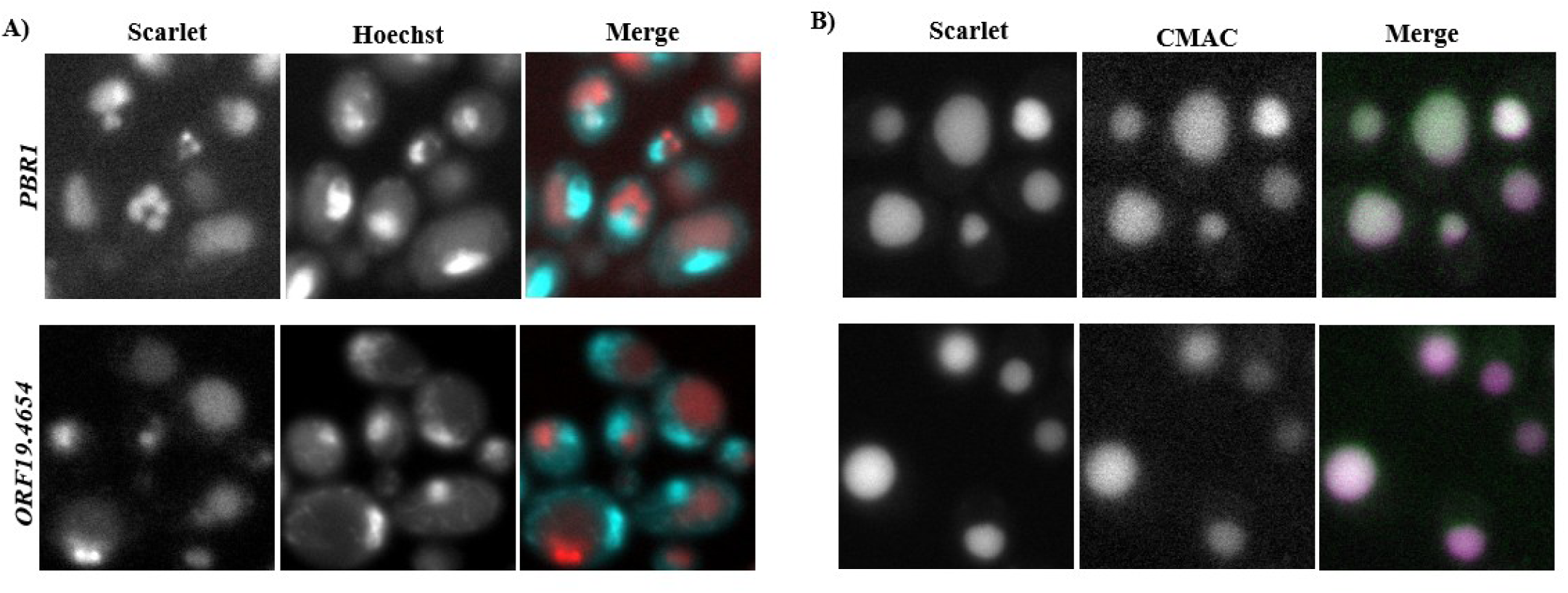
Pbr1 and Orf19.4654 localize in the vacuole. (A) *PBR1* and *ORF19.*4654 stained with 1µg/ml Hoechst 33342 for nuclear localization. Scarlet and Hoechst signals show different localizations in the cell. (B) *PBR1* and *ORF19.*4654 were stained with 10 mM CMAC for vacuolar localization. Scarlet and CMAC signals show the same localizations in the cell. First row *PBR1*, second row *ORF19.*4654.

### Colocalization analysis suggests that Ppp1 is not part of Sur7-defined eisosomes

Due to the punctate surface distribution of Ppp-1-GFP in biofilm yeast cells, we hypothesized that it may be a component of the eisosome, a subcellular domain of the plasma membrane involved in endocytosis and stress responses (Douglas et al. 2011). Repeated co-occurrence of two probes within a cell is an indicator of the presence of the two proteins in the same compartment. We constructed a double tag with Sur7, a marker of the eisosome (Dunn et al., 2011), and assessed protein distribution. As seen in Figure 4A, although the GFP and Scarlet signals for Ppp1-GFP and Sur7-Scarlet show some overlap, a lot of the GFP signal remained distinct. We measured this using Manders’ and Pearson’s Correlation Coefficient to quantify this puncta colocalization, as these measure the fractional overlap of the intensity of one signal with the other, and the correlation between signal intensities in the channels (Dunn et al., 2011).

**Figure 4:**
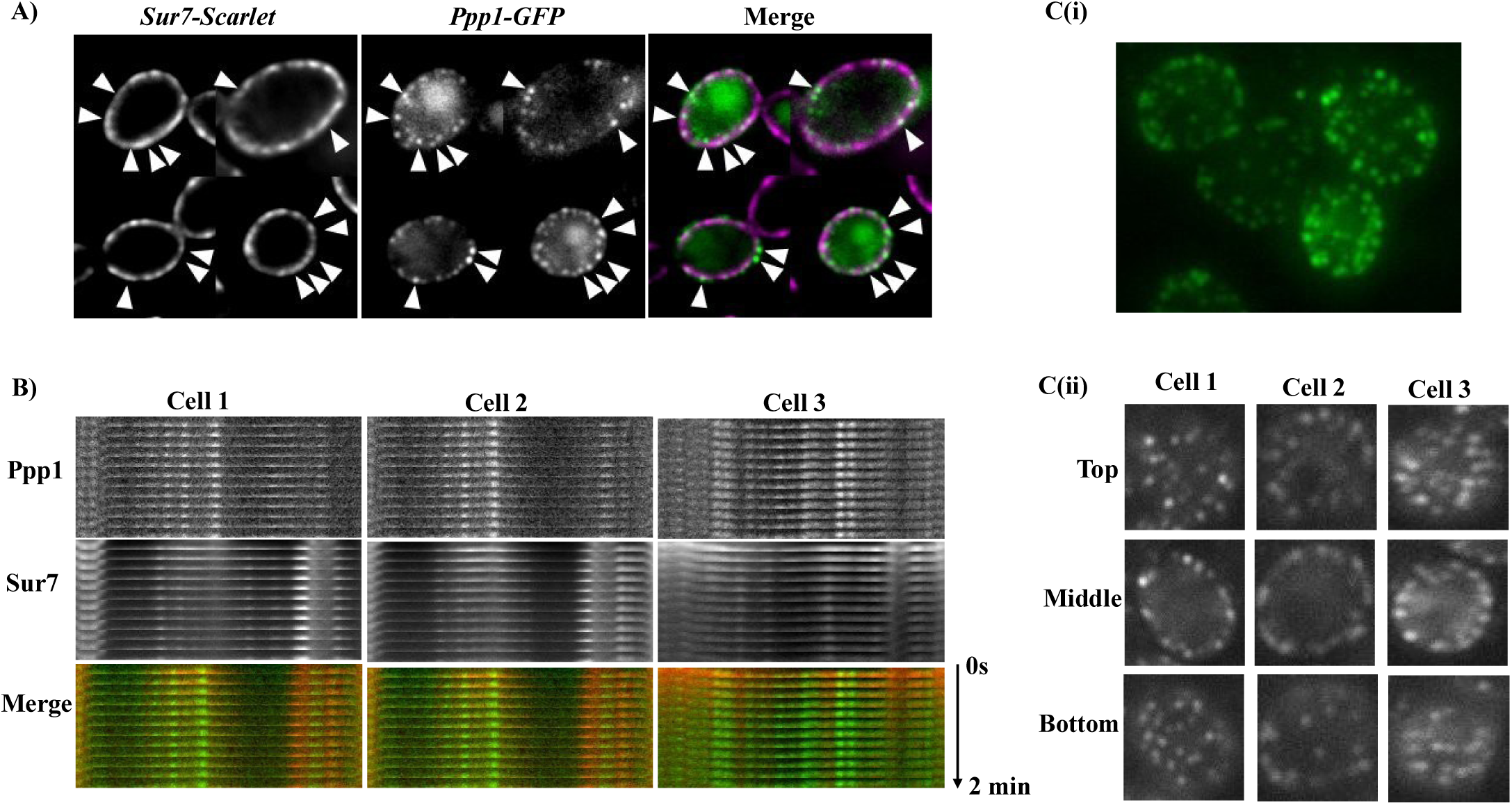
Fluorescence microscopy screen for subcellular localization of Ppp1. (A) Colocalization between Ppp1 visualized by merging the GFP and Scarlet channels. Arrows highlight puncta. (B) Kymogram for further puncta colocalization between Sur7 and Ppp1. Images were captured at 10s intervals for 2 mins. The horizontal and vertical arrows indicate puncta distribution and time, respectively. (C) i) Orthogonal views of Ppp1 distribution demonstrating peripheral puncta. ii) Z-stacks of Ppp1 outlining puncta distribution throughout the cell.

According to Mander’s Correlation Coefficient, the proportion of Ppp1-positive pixels that overlap with Sur7-Scarlet pixels is 0.45 ± 0.11 (N=40), and the correlation between signal intensities is 0.36 ± 0.09 (N=40), demonstrating that these signals overlap partially. Examination of the distribution of the punctate protein Cwr1 reveals that this protein appears to be motile in the plasma membrane and transiently associated with Sur7 (Naseem et al., 2024), suggesting that Ppp1 may also engage in similar behaviour. To assess this, we performed time-lapse imaging of Sur7-Scarlet and Ppp1-GFP. Images were captured of both proteins at 10s intervals for 2 min to generate kymographs to compare puncta localization and motility. As seen in Figure 4C, both Sur7 and Ppp1 kymogram signals are stable and non-overlapping, indicating that Ppp1 shows partial co-localization with the Sur7 component of the eisosome and is not likely to be located in the eisosome, even transiently.

### Deletion of *PPP1* does not affect biofilm structure

To examine the overall biofilm structure when *PPP1* is deleted and visualize possible changes, WT, *ppp1Δ/Δ,* and *ppp1Δ/Δ* + *PPP1* strains were grown in RPMI with or without 10% FBS for 24 hours. No changes were observed between the WT and the mutant strain (Figure 5A). A minor difference in biofilm formation was observed in the complement strain. In the apical view for growth in RPMI, while the *ppp1Δ/Δ* strain formed a biofilm similar to the WT strain, the complemented *ppp1Δ/Δ* + *PPP1* strain formed a somewhat more extensive biofilm than either the WT or the mutant strain (Figure 5A). When 10% FBS was added (Figure 5B), as expected, overall biofilm formation increased, but now the complemented strain had both shorter hyphae and produced less biofilm compared to the mutant and WT. Deletion of *PPP1* does not have an observable effect on biofilm formation, but reinsertion of *PPP1* at its native locus causes variable morphological changes in biofilm structure.

**Figure 5:**
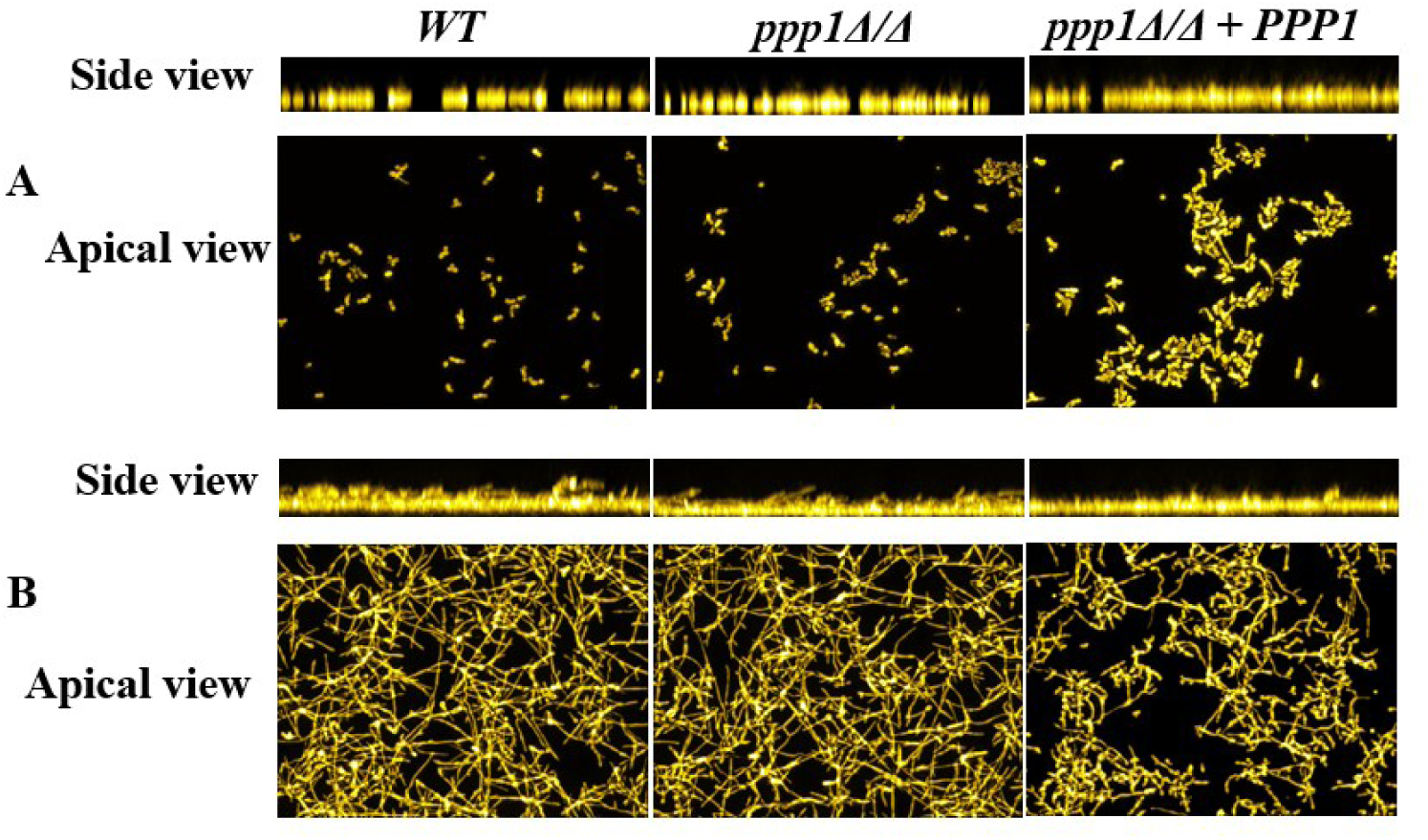
Optical sectioning shows that *ppp1Δ/Δ+PPP1* does not behave like the WT. Side and apical view confocal microscopy images of WT, mutant (*ppp1Δ/Δ*), and complemented strain (*ppp1Δ/Δ+PPP1*) grown for 24 hours at 37^0^C in **A) RPMI, B) RPMI + 10% FBS**. *ppp1Δ/Δ* behaves like the WT under both media. However, *ppp1Δ/Δ+PPP1* forms longer hyphae in RPMI and less biofilm in RPMI + 10% FBS. Imaging was done in duplicates.

### Deletion of *PPP1* does not impact biofilm biomass

We measured biofilm biomass in the mutant and complemented strains. In buffered Spider medium, deletion of *PPP1* did not cause any difference in biofilm production, as the biomass of all strains was similar (Figure 6). Biofilm formation in unbuffered Spider media biofilm production was also low as the buffered media. Under these conditions, the mutant produced a bit more biofilm than the WT with OD595 = 0.75 (7.5 × 10^6^ cells) but less than the complemented strain of 9 × 10^6^ cells. There was a 2.5-fold increase when cells were grown in RPMI media. Here, there was a reduction in biomass in the mutant strain (1.8 × 10^7^ cells) compared to the complemented strain, which produced almost the same amount of biofilm (2.2 × 10^7^ cells) as the WT (2.4 ×10^7^ cells). This data suggests that the biofilm biomass is not impacted by the loss of *PPP1,* but instead by the type of media in which the cells are cultured.

**Figure 6:**
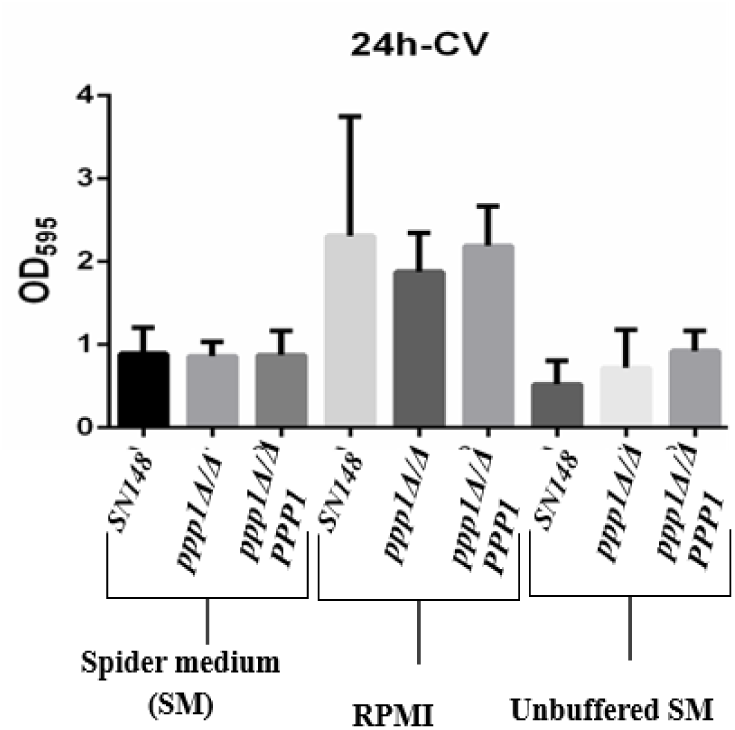
Crystal violet shows that Ppp1 does not impact biofilm biomass. Crystal violet assay measuring the biofilm biomass of *ppp1Δ/Δ* and *ppp1Δ/Δ +PPP1,* comparing it to the *SN148 (WT)*. Experiments were done in triplicate and repeated for consistency. Error bars represent standard deviation.

### *PPP1* is not involved in stress and antifungal response

*C. albicans* has an environmental response pathway made up of genes that respond to factors like osmotic and oxidative stress (Ruis & Schüller, 1995). We tested these stress factors at 30^0^C and 37^0^C on the mutant (*ppp1Δ/Δ*) and complemented strain (*ppp1Δ/Δ* + *PPP1*) to see if loss of *PPP1* is affecting response to stress inducers and/or any of the classes of antifungals. To identify distinct phenotypes in the stress assays, concentrations at which cells can grow at varying rates and morphologies were optimized from literature review (Rashid et al., 2022). No altered growth was identified in any of the stress indicators as seen in Figure S2A, suggesting that Ppp1 is not important for osmotolerance or oxidative stress.

The effect of Ppp1 on antifungal resistance was evaluated for the four classes of antifungals to determine if Ppp1 has a specific role in pathways and subcellular compartments associated with these antifungals. Caspofungin, an echinocandin, inhibits 1-3 β-D-glucan affecting the cell wall synthesis; amphotericin B, a polyene, integrates into the ergosterol membrane of Candida; fluconazole, an azole, inhibits cytochrome P450-dependent 14α-lanosterol demethylase inhibiting ergosterol biosynthesis, and 5-flucytosine (5-FC), a pyrimidine analog, disrupts DNA and RNA synthesis (Taff et al., 2013, Perfect, 2017). As with the stress assays, we did not observe any altered growth in the mutant and complemented strain in comparison with the wild type (WT) (see Figure S2B).

## DISCUSSION

Biofilms are a virulence factor of medical importance due to their prevalence in nosocomial infections, especially among patients with implanted devices (Wisplinghoff et al. 2003, Atriwal et al., 2021). They are also a leading component of infections in immunocompromised patients and can affect healthy individuals. Due to the nature of biofilm cells, they are resistant to most antifungals, and they can contribute to systemic candidiasis that can lead to 30-50% mortality rates in infected patients (Atriwal et al., 2021).

Decades of research have gone into understanding how *C. albicans* biofilms are formed and into defining the network of genes and transcription factors involved in this process. Nobile et al. 2012 identified more than 1000 genes involved in biofilm formation, many of which have not been investigated, and some of which are unique to *C. albicans* and its closest relatives. Uncharacterized genes that are highly induced under biofilm conditions represent an interesting target for further investigation due to the role of biofilms in antifungal resistance. We selected, for further investigation, three poorly characterized, highly upregulated genes unique to *C. albicans* and close relatives that encode small proteins with N-signal peptides, namely *PPP1*, *PBR1,* and *ORF19.4654* (Nobile et al., 2012).

Microscopic investigation into the subcellular localization of fluorescently tagged proteins can provide insights into protein function and cellular physiology (Combs & Shroff, 2017). This technique has been used in numerous studies defining protein subcellular localization in fungi (Augstein et al., 2003; Chang et al., 2023), and we have applied it to the investigation of the proteins encoded by these biofilm-induced genes.

The fluorescently tagged Pbr1 and Orf19.4654 showed similar protein distribution under both planktonic and biofilm conditions, and both proteins were localized to the vacuole. Even though the RNA Seq data showed that the expression of the genes was dramatically induced in biofilm cells relative to planktonic cells, the protein concentration in the vacuole appeared similar in both cell types. Vacuolar localization is a frequent consequence of signal-peptide-directed protein sorting, and thus may define the proper subcellular localization of both Pbr1 and Orf19.4654. The vacuole is part of the secretory pathway involved in cellular regulation and often utilizes signal peptide sorting to direct proteins across the secretory pathway and to other subcellular organelles. Proteins like Cpy1 and Apr1 have signal sequences directing them to the vacuole, similar to the Kex proteins localized in the Golgi (Mukhtar et al., 1992, Niimi et al., 2006).

An alternative hypothesis posits that the localization of Pbr1 and Orf19.4654 arises from protein misfolding due to the added fluorescent tag, leading to the misfolded proteins being directed to the vacuole for degradation. Studies have shown how misfolded/damaged proteins that successfully move through the secretory pathway, evading the ER-associated degradation (ERAD) pathways, multi–vesicular body (MVB) pathway, and other degradation mechanisms are lysed in the vacuole (Wang et al., 2011; Wang & Ng, 2010). However, Pbr1 and Orf19.4654 were constructed with a linker between the protein and fluorescent tag to prevent misfolding and improve FP stability (Chen et al., 2013). Studies have shown that these linkers can even improve protein function (Sabourin et al., 2007; Werner et al., 2006).

The other protein, Ppp1, showed a dramatically different localization from Pbr1 and Orf19.4654. This protein showed distinct puncta only on the cell periphery of yeast cells grown under biofilm-inducing conditions and a weak, non-localized signal in planktonic cells. Similar to Pbr1 and Orf19.4654, the hyphal cells that formed under the biofilm conditions also had a non-localized, low level of the Ppp1-GFP signal.

Due to the protein localization pattern of Ppp1, colocalization with Sur7, an eisosome marker with similar distribution, was investigated to further define its subcellular location. Puncta distribution at the cell surface for Sur7 is the same for yeast and hyphae cells, unlike Ppp1, where the puncta distribution is mostly limited to yeast-form cells. Merged images suggest infrequent co-localization of Ppp1 and Sur7. Higher resolution colocalization qualification with kymograms shows that although appearing similar in size and location, the puncta of Ppp1 and Sur7 essentially localize separately, implying that Ppp1 is not part of the Sur7-defined eisosome. Both proteins remain immobile throughout the 2-min time lapse, suggesting they may belong to different cell surface protein complexes. A recent study (Lanze et al., 2025) was conducted to identify novel eisosome proteins using a proximity labelling technique, TurboID. Sur7 was used as a bait, and a number of new and previously poorly characterized proteins were identified as interacting partners. Ppp1 was not found as part of these Sur7-interacting proteins, consistent with our observation that Ppp1 does not localize directly with Sur7.

Biofilm assays of the null mutant and complemented reinsertion showed that, although *PPP1* was highly upregulated in biofilms, loss of Ppp1 had little impact on biofilm formation. The null mutation resulted in no apparent change in biofilm mass measured in the crystal violet assay. As well, fluorescence microscopy to determine the biofilm structure found little difference in the biofilms generated by the WT and null mutant strains. However, this assay detected some minor differences in the complemented strain relative to the null mutant and the WT, as somewhat more extensive biofilms formed in RPMI growth medium, and fewer biofilms were generated when FBS was added to the biofilm induction medium.

A variety of other assays did not identify phenotypic consequences of *PPP1* deletion. Environmental factors such as stresses can impact biofilm virulence, so we evaluated the effect of NaCl, CaCl_2_, glycerol, and H_2_O_2_ (Pemmaraju et al., 2016). Annotation by phenotypic analysis of mutant strains is a powerful, standard technique for functional characterization of proteins (Winzeler et al., 1999). This can be used to identify drug targets and study the effect of different stress responses on the cell (Wagner, 2000). The absence of altered growth when cells are treated with stress factors suggests that Ppp1 is unlikely to be involved in stress responses in *C. albicans*. Drug resistance is a major area of medical research due to the high mortality rate associated with Candida infections. Screening the *PPP1* deletion strain with fluconazole, 5-flucytosine, caspofungin, and amphotericin B, representing the four classes of antifungals, shows no interaction between Ppp1 and the antifungals.

There are other complexes/subcellular domains with similar cell periphery puncta distribution as Ppp1, but show characteristics that suggest they are distinct from Ppp1. The plasma membrane domains, MCP (Pma1 domain), and the TORC2 complex both form distinct cell surface puncta (Douglas et al., 2011; Malinsky et al., 2010). However, unlike the stable Ppp1 puncta, the TORC2 complex forms a module showing dynamic puncta movement (Malinsky et al., 2010; Berchtold & Walther, 2009). As well, while Pma1-defined puncta are stable, they have a much higher density than the Ppp1 puncta, with Pma1 being the most abundant membrane protein (Malínská et al., 2003; Malinsky et al., 2010). If Ppp1 were associated with Pma1, it would be with a small subset of the complexes. Other plasma membrane proteins not localized in a sub-domain have a more evenly distributed punctate pattern than the domain proteins (Bean et al., 2022; Malínská et al., 2003). Investigating the relationship between Ppp1 and the Pma1 domain can support Ppp1’s subcellular localization and provide more insight into its functions.

## MATERIALS AND METHODS

### CRISPR constructs and strains

The *C. albicans* strains are described in Supplemental Table 1. The complemented strain *ppp1Δ/Δ + PPP1* (*orf19.7608Δ/Δ*+ *ORF19.7608*) was constructed as described by Costa et al., 2020, and all other strains were constructed using the CRISPR-Cas9 transient method (Min et al., 2016). In the transient method, Cas9, sgRNA, and repair DNA are amplified from plasmids using PCR and transformed into the appropriate background strain. All constructs were identified by colony PCR using external and internal primers. All primers used are listed in Supplemental Table 2.

The mutant strain *ppp1Δ/Δ* (*orf19.7608Δ/Δ*) was constructed by deleting both copies of the gene in SN148. Cas9 was PCR amplified from pV1093, while the repair DNA was amplified from pFA-Arg4. The repair primers contained about 80 bp of homology to the 5’ and 3’ ends of the non-coding region of the deletion gene. An sgRNA with the best on-target/off-target scores was chosen from the genomic region using Benchling to increase the efficiency of the target cut. DNA amplification was conducted using Onetaq 2x MM with standard buffer (New England Biolabs). The complement strain *ppp1Δ/Δ + PPP1* (*orf19.7608Δ/Δ*+ *ORF19.7608*) was constructed by reintroducing both alleles of *PPP1* (*ORF19.7608*) into the mutant strain *ppp1Δ/Δ* (*orf19.7608Δ/Δ*). This was done using the conventional CRISPR-Cas9 method: the pV1093-gRNA *ARG4* plasmid was integrated into the DNA (*ENO1*), and the selectable marker *ARG4* was replaced on both alleles by *PPP1* (*orf19.7608Δ/Δ*).

To construct the tagged strains, the fluorescent protein (FP) and selectable marker were inserted to replace the stop codon and about 150-200bp downstream of the gene (Huh et. al, 2003). This allowed integration of the FP without disruption of protein expression of the gene of interest. For *PPP1-GFP*, the repair and selectable marker were amplified from pFA-GFP-HIS1. The CIP10-mScarlet-I IDT plasmid used to amplify Scarlet does not have a selection marker, so for *PBR1* and *ORF19.4654*, the repair DNAs were constructed as cassettes made up of Scarlet from CIP10-mScarlet-I IDT and *URA3* from pFA-GFP-CaUra3. This was done in three rounds of PCR, similar to the sgRNA cassette, with the first round involving amplifying *URA3* from pFA-GFP-CaUra3 and Scarlet from CIP10-mScarlet-I IDT. These amplicons form a single joint template in the second round of PCR. In the third round, homology arms complementary to the 3’ end of the genes and the downstream non-coding region were introduced. During this third round, a linker was introduced through the forward repair primer to further aid the proper fusion of the protein. Two sgRNAs were used to improve CRISPR efficiency (Zhang et al. 2022, Chen et al. 2014).

### Yeast transformation

The transformations were done using a modified Lithium acetate method (Walther and Wendland 2003). First, cells were grown overnight at 30^0^C in YPD with uridine (10 g yeast extract, 20 g peptone, 20 g glucose, 50mg uridine, 1l water), shaking at 220 rpm. They were then grown to log phase (OD_600_=0.8) to improve transformation competency. 10 ml of culture was centrifuged, washed twice with nuclease-free water, and incubated in 600µl 1x LiAc for 20 mins to improve the cells’ ability to absorb foreign DNA. In an Eppendorf tube, 100µl of the competent Candida cells, 10µl of single-stranded salmon sperm DNA (heated for 10 mins at 98^0^C), 1µg CaCas9, 1µg sgRNA, and 3-6µg repair DNA were added. 600µl 1xLiAc/ 40% PEG was also added to the mix, and this was incubated for 3 hours at 30^0^C. Next, the transformation mix was put in a water bath at 44^0^C for 15 mins and then on ice for 1 min. The mix was then centrifuged for 10 mins at 6000 rpm, and the pellet was washed with nuclease-free water. The pellet was then resuspended in 900µl YPD with uridine and grown for 2 hours to allow cell recovery. Another centrifugation was done to remove the media. Cells were washed and resuspended in nuclease-free water and then plated on selection plates and grown at 30^0^C for 3-4 days.

### Biofilm characterization

To assay biofilm formation in a 96-well plate (Greiner 96-well plate; catalog no. 655090), strains were inoculated to an optical density at 600 nm (OD600) of 0.05 from overnight cultures into 100μl of prewarmed 0.2% glucose containing RPMI with or without 10% fetal bovine serum (FBS). First, the cells were incubated in a shaker incubator at 37^0^C for 90 mins with mild shaking (60 rpm) to allow adherence, and then each well was gently washed twice with 1x PBS. Next, 100μl of prewarmed RPMI with or without 10% FBS was added into each well, and cells were allowed to form biofilms in a shaker incubator with 60 rpm at 37^0^C for 24 hours. At that time, the medium was discarded from each well, and biofilms were fixed by incubation with 100μl of 4% formaldehyde in 1x PBS for 1 hour and then gently washed twice with 1x PBS. Biofilms were stained with calcofluor white (200μg/ml in PBS) overnight at room temperature (RT) with mild shaking (60 rpm), and then each well was gently washed twice with 1x PBS. To clarify biofilms in the 96-well plates, we used 100% TDE (thiodiethanol), which has a refractive index of 1.521. The PBS was removed from each well and replaced with 100μl of 50% TDE in PBS. Cells were incubated in 96-well plates at room temperature for an hour, and then the solution was removed from each well. Finally, 100μl of 100% TDE solution was added to the wells and incubated at RT for an hour, when the clarified biofilm was transparent. The biofilms for strains of *WT*, *ppp1Δ/Δ,* and *ppp1Δ/Δ + PPP1 (WT, orf19.7608Δ/Δ*, *orf19.7608Δ/Δ*+ *ORF19.*7608) were imaged using a Keyence BZ-X800E fluorescence microscope (Sharma et al. 2023).

Optical sections of the biofilms were collected in several series of planes at a 0.45- or 0.85-μm step size. The stacks were concatenated and processed using Fiji (Schindelin et al. 2012). The images were processed using the Background Subtract plugin. The orthogonal projection images were obtained by reslicing the stack and using the maximum intensity Z-projection. The scale of the stacked side-view images was adjusted based on the objective used for the imaging.

### Crystal violet assay

Biomass accumulation in biofilms was measured by the crystal violet assay. *WT*, *ppp1Δ/Δ,* and *ppp1Δ/Δ + PPP1* (*WT, orf19.7608Δ/Δ*, *orf19.7608Δ/Δ*+ *ORF19.7608*) were cultured in YPD with uridine overnight at 30^0^C /220 rpm. Cells were washed twice with 1x PBS and resuspended in Spider media. The concentration of cells was adjusted to OD600 = 0.5, and 200µl of the inoculum was transferred to a 96-well plate with a non-treated surface for initial incubation at 37^0^C for 1.5 h in static conditions to allow cell adherence. Next, non-adherent cells were washed off with 1x PBS, and 200 µl of either unbuffered spider, RPMI, or buffered spider media was added to each well. Strains were grown in different biofilm-inducing media to identify potential differences in biofilm biomass based on the chemical composition of these media. The plates were incubated for 24 h at 37^0^C/75 rpm in the shaking incubator. The biofilms were washed once with 1x PBS, air dried for 45 mins, and stained with 0.4% aqueous crystal violet solution for 45 min at room temperature (RT). Next, the biofilms are washed twice with autoclaved distilled water and then de-stained with 95% ethanol for 45 mins. The de-stained solution was diluted at 1:10 in 95% ethanol, and the absorbance was measured at OD_595_ (Tecan Infinite M200 Pro, Grodig, Austria). The procedures were performed in 6 wells for each test.

### Cellular stress assay

The stress response of *ppp1* mutant strains was analyzed by growing cells in YPD uridine under different stress conditions. Single colonies of the WT, mutant, and complemented strains (*WT, ppp1Δ/Δ,* and *ppp1Δ/Δ + PPP1*) were grown overnight in YPD with uridine at 220 rpm, 30^0^C. Cells were washed twice in 1x PBS, and calculations were made for OD600 = 1×10^7^ and serial dilutions to 10^3^. 3µl of samples were spotted on agar plates and grown for two days (except caspofungin plates, where growth was for 5 days) (Rashid et al. 2022). Strains were subjected to osmotic stress of CaCl_2_ (400 mM), NaCl (500 mM), glycerol (100 mM), and oxidative stress of hydrogen peroxide (3 mM). Antifungal resistance to the different classes of antifungals: fluconazole (1µg/ml), 5-flucytosine (0.5µg/ml), amphotericin B (0.25µg/ml), and caspofungin (0.75µg/ml) was also determined. Growth at 30^0^C and 37^0^C analyzed the effect of temperature on the strains. Experiments were repeated three times for each test to evaluate reproducibility. The selected concentrations of salts, glycerol, hydrogen peroxide, and antifungals were determined from literature reviews (Galdiero et al., 2020, Ejaz & Ramzan, 2019, Roscetto et al., 2018, Rashid et al., 2022). Images were captured using an Epson Perfection v500 photo scanner.

### Subcellular fluorescence microscopy

Images of Orf19.7608 (Ppp1), Pbr1, and Orf19.4654 were captured with an epifluorescence microscope (Schindelin et al. 2012). The three biofilm genes were grown under both biofilm and planktonic conditions to visualize their protein distribution and identify any difference in pattern under these conditions. For planktonic growth, cells were grown in Falcon tubes at 220 rpm, 30^0^C for 12-16 hours in YPD media supplemented with uridine. Biofilm growth was assessed in 6-well plates incubated at 37^0^C for 48 hours using Spider media.

Nuclear dye Hoechst 33342 (1μg/ml) and vacuolar dye CMAC (10 mM) were also used as co-labels to query the subcellular distributions of Pbr1 and Orf19.4654. Cells were grown overnight in YPD, washed twice with 1x PBS, stained for 20 mins in the dark, washed twice in 1x PBS to remove excess dye, and visualized. Cells were imaged on a Leica DMi6000 microscope with 100x Plan Fluotar lens (NA 1.3), Hamamatsu Orca CCD camera, Leica EL600 Hg light source, and appropriate filters for Hoechst/CMAC (385/20x, 445/70m), eGFP (472/30x,520/35m), and Scarlet (562/40x, 624/40m).

Colocalization was analyzed between the GFP and Scarlet in Ppp1-GFP Sur7-Scarlet expressing cells using the BIOP implementation of the JaCoP plugin (Bolte & Cordelières, 2006). Kymograms of Ppp1-GFP/Sur7-Scarlet expressing cells were produced by capturing single-slice epifluorescence images of both channels every 10 seconds for 2 minutes. A Fiji script was run by defining the outline of the cell as an elliptical selection; this was converted to a straight line with a width of 10 pixels, and the resulting images were arranged into the kymograph stacks seen in Figure 4B.

## Acknowledgments

This project was funded by NSERC-CREATE and SynBioApps grants. We appreciate the Centre for Microscopy and Cellular Imaging, Concordia University, for providing us with the microscopes and technical capacity to visualize and analyze images.

