## Supplementary Data for "Characterization of *ORF19.7608* (*PPP1*), a Biofilm-induced Gene of *Candida albicans*"

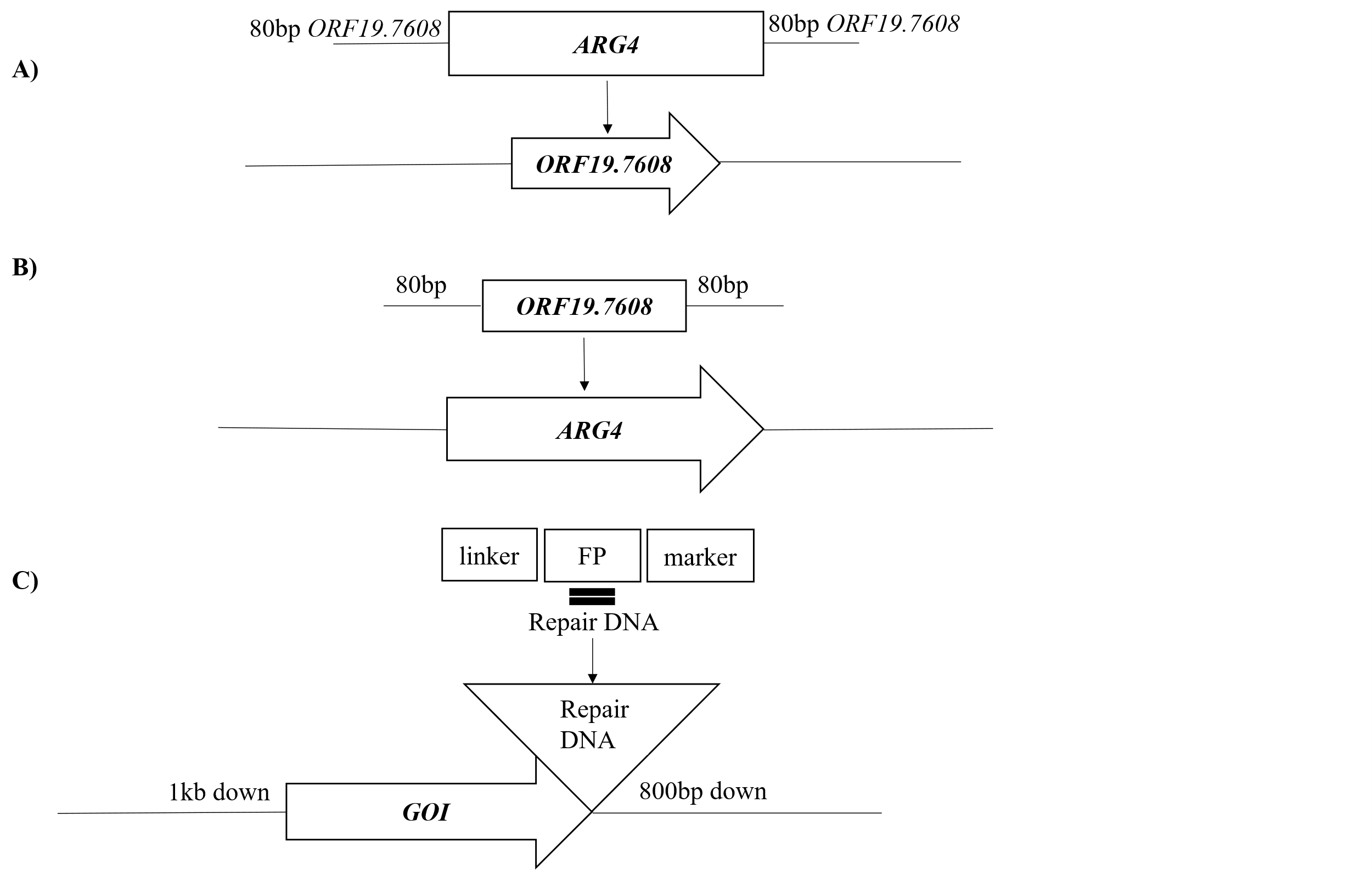

**Figure S1: Schematic representation of the construction of mutant, complement, and tagged strains**

A) The mutant of *ORF19.7608* was constructed by replacing both alleles of *ORF19.7608* with *ARG4*. The *ARG4* repair DNA contained 80bp homology arms to *ORF19.7608,* ensuring homologous recombination.
B) The complement strain was constructed by replacing *ARG4* in the mutant strain with *ORF19.7608*. The repair DNA was constructed with 80bp homologous arms to *ORF19.7608*, ensuring homologous recombination.
C) Tagged strains were designed as C-terminal fusion constructs with repair DNA replacing the stop codon and 200bp downstream sequence.

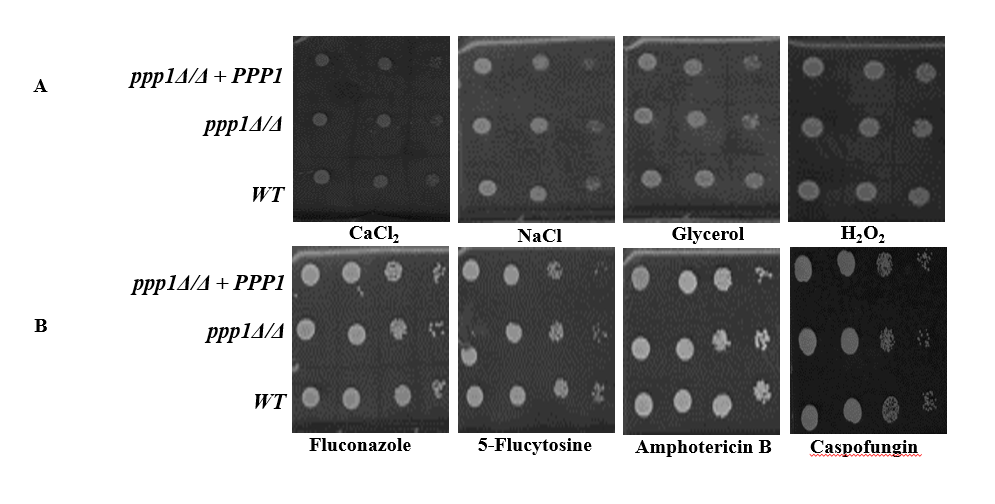

**Figure S2: Deletion of *PPP1* does not impact stress responses and antifungal resistance**

1:10 serial dilution of overnight cultures grown in yeast growth conditions was spotted (highest cell concentrations being 1 x 10^6^ cells/ml and lowest being 1 x 10^4^ cells/ml or 1 x 10^3^ cells/ml) onto YPD agar plates containing different chemicals and incubated at 30^o^C for 2 days except for caspofungin (5 days).

(A) Strains were subjected to osmotic stress of CaCl2 (400mM), NaCl (500mM), glycerol (100mM) and oxidative stress of H_2_O_2_ (3mM)

(B) Determination of resistance to different antifungal drugs: fluconazole (1µg/ml), 5-Flucytosine (0.5µg/ml), amphotericin B (0.25µg/ml), and caspofungin (0.75µg/ml).

Experiments were repeated three times for each sample for consistent results. Results were the same at 30^0^C and 37^0^C

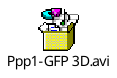

**Supplemental Video 1**: **Orthogonal view of Ppp1 showing puncta distribution in 3D**

**Supplemental Table 1: Strains Used in this Study**

| **Strains** | **Parent** | **Description** |
| --- | --- | --- |
| *ppp1Δ/Δ* | *SN148 a/α* | *orf19.7608*Δ::*ARG4*/*orf19.7608*Δ::*ARG4* |
| *ppp1Δ /Δ* + *PPP1* | *ppp1Δ/Δ* | *ORF19.7608*/*ORF19.7608*; *his1*/*his1*; leu2/leu2; *arg4/arg4*; *ura3*::imm434/*ura3*::imm434; *iro1*::imm434/*iro1*::imm434 + pV1093-gRNA *ARG4* |
| *PPP1-GFP* | *SN148a/α* | *ORF19.7608*-*GFP*-::*HIS1*/*ORF19.7608-GFP::HIS1*; *leu2/leu2; arg4/arg4;* *ura3*::imm434/*ura3*::imm434;*iro1*::imm434/*iro1*::imm434 |
| *SUR7-Scarlet PPP1-GFP* | *PPP1-GFP* | *ORF19.7608-GFP-::HIS1/ORF19.7608-GFP::HIS1*; *SUR7-Scarlet::URA3/ SUR7-Scarlet::URA3; arg4/arg4, leu2/leu2* |
| *PBR1-Scarlet*  *PPP1-GFP* | *PPP1-GFP* | *ORF19.7608-GFP-::HIS1/ORF19.7608-GFP::HIS1; PBR1-Scarlet::URA3/ PBR1-Scarlet::URA3; arg4/arg4, leu2/leu2* |
| *ORF19.4654-Scarlet* | *SN148a/α* | *ORF19.4654-Scarlet::URA3/ ORF19.4654-Scarlet::URA3; arg4/arg4, leu2/leu2; his1/his1* |

**Supplemental Table 2:** **Oligonucleotides Used in the Study**

| **Name** | **Sequence (5’ to 3’)** |
| --- | --- |
| ***ppp1Δ/Δ*_sgRNA_F** | GATGACGGTGATGATGATGAgttttagagctagaaatagcaagttaaa |
| ***ppp1Δ/Δ*_sgRNA_R** | TCATCATCATCACCGTCATCcaaattaaaaatagtttacgcaagtc |
| ***ppp1Δ/Δ*_repair_F** | TCATTAGCAAAGCAGTTGTAACAAACAAACAAAAAGACAACAATAATAAAAATCCATATCACACAATTACAATAAATCgaagcttcgtacgctgcaggtc |
| ***ppp1Δ/Δ*_repair_R** | TGAAATAAAACAAAAATAAAACAACACAAAGTTGAATAAGAAGAAGAAAGATTCAAATAGTACGGCTTGGCTATGTtctgatatcatcgatgaattcgag |
| ***ppp1Δ/Δ*_diag_F** | GGAGTCGTCAACTGCAAATTGTGAAG |
| ***ppp1Δ/Δ*_diag_F** | ATTAGCCAAGGCAGGATGTATAGC |
| ***ppp1Δ/Δ+PPP1*-FW** | gtcgacTGACTGTTTGTTTCAATTC |
| ***ppp1Δ/Δ+PPP1*-Rv** | acgcgtGCAGATAGTGGCATG |
| **Fw_Int_*ppp1Δ/Δ+PPP1*** | GAAGACTTCAGTCGTTTTAGCTGC |
| **RPF1-R** | CGCCAAAGAGTTTCCCCTATTATC |
| **RPF-1** | GAGCAGTGTACACACACACATCTTG |
| **Diag_RP10_F** | CATGAGGCCTCCATGAGGCCTC |
| **Diag_CIp10_R** | agctatgaccatgattacgccaagc |
| ***URA3*-F** | ggagttggattagatgataaaggtgatgg |
| ***PPP1*-*GFP*_sgRNA_F** | GTAGTGAATATATTTGAAAAgttttagagctagaaatagcaagttaaa |
| ***PPP1*-*GFP*_sgRNA_R** | TTTTCAAATATATTCACTACcaaattaaaaatagtttacgcaagtc |
| ***PPP1-GFP*_repair_F** | GAGATTACTTTTCTCAATGGAAACAAGGACTCGACAACTTAATTCAAAAAGGTAAGACATGGTTTAGTGGTCTTTTCGGTggtgctggcgcaggtgcttc |
| ***PPP1-GFP*_repair_R** | ATATTATGAGAAAAAAAATTTGATATACCTAAGAATTGAAACCTGTAAATGACAAAAAATTTATAAACAAATAAAAATCCccgcataggccactagtgga |
| ***PPP1-GFP*_Diag_Int_F** | GAAGACTTCAGTCGTTTTAGCTGC |
| **Con_*PPP1*-R1** | ATTAGCCAAGGCAGGATGTATAGC |
| ***PPP1-GFP*_Diag_Ext_F** | TGAAGCACCTTCTTTACAGCAACACG |
| ***GFP*_R** | tctttcgaaagggcagattgtgtgg |
| ***HIS1*_F** | gcagatggcgagtacgaaaagc |
| ***PPP1-GFP*_Diag_Ext_R** | CAACTTTCACAAGTGCAGATAGTGGCA |
| ***SUR7-*Scarlet_sgRNA_F** | CGTATATTAAATATACCAATgttttagagctagaaatagcaagttaaa |
| ***SUR7-Scarlet*_sgRNA_R** | ATTGGTATATTTAATATACGcaaattaaaaatagtttacgcaagtc |
| ***SUR7-Scarlet*_repair_F** | CAGGCGGTATTAGATTCTTCAAAATCAAAAGAAACCAAAAAGTTTCCGATGATGAATCAGTAggtgctggcgcaggtgctatggtcagtaaaggggaagc |
| ***SUR7*-Scarlet_repair_R** | TAAAGATTCCAATAATGGTAATACTGATAATAATAATAATAATAACAATGATGATTTTAGTAATAGTAGTAACAGTtctgatatcatcgatgaattcgag |
| ***SUR7-Scarle*t_diag_ext_F** | GTCTCATTGCCCTTGCATTCAGTG |
| ***SUR7*-*Scarlet*_diag_ext_R** | CACAATCCATATGTAACTCATGTCACG |
| **Diag_*Scarlet*_R2** | GGCATTTGCACAGGTTTCTTAGC |
| **Diag_*URA3*-F1** | gaaactcatgcctcaccagtagc |
| ***ORF19.4654-Scarlet*_sgRNA1_F** | CTTCTCTGTACATTAAGTTAgttttagagctagaaatagcaagttaaa |
| ***ORF19.4654-Scarlet*_sgRNA1_R** | TAACTTAATGTACAGAGAAGcaaattaaaaatagtttacgcaagtc |
| ***ORF19.4654-Scarlet*_sgRNA2_F** | CTACTCATTGGTTGTTCTAGgttttagagctagaaatagcaagttaaa |
| ***ORF19.4654-Scarlet*_sgRNA2_R** | CTAGAACAACCAATGAGTAGcaaattaaaaatagtttacgcaagtc |
| ***ORF19.4654-Scarlet*_repair_F** | ATATTCAATGGAAATTATTATTAGGTTGTTTTTTATTTGCAATTGTAAGTTTACTTGCAATGggtgctggcgcaggtgctatggtcagtaaaggggaagc |
| ***ORF19.4654-Scarlet*_repair_R** | TAGAACTGAATCTGGGTATATTTTTTGTTAATTTCGGTCCGATTTAGCTATTTGCTATGGTATTAAGTTCCCCATTtctgatatcatcgatgaattcgag |
| ***ORF19.4654-Scarlet*_diag_ext_F** | CAGCAACAACATTGTCAACCACCAA |
| ***ORF19.4654-Scarlet*_diag_ext_R** | CGCGCGTTCTTTTCTCCTCCT |
| ***PBR1-Scarlet*_sgRNA_F** | GTTTAATTAATTGTTAAGTTgttttagagctagaaatagcaagttaaa |
| ***PBR1-Scarlet*_sgRNA_R** | AACTTAACAATTAATTAAACcaaattaaaaatagtttacgcaagtc |
| ***PBR1-Scarlet*_repair_F** | ATCTGCCACCAAAGAACTCCTACAAAGTTAGTATCTACGGTCGTCTCAATTGGGCTGTCTTGggtgctggcgcaggtgctatggtcagtaaaggggaagc |
| ***PBR1-Scarlet*_repair_R** | TTATATATATATGTACACATAGTATAAGAACTAATAAATAAGCAACGAGAAGAAACGAAATAAAACAACGACAACGtctgatatcatcgatgaattcgag |
| ***PBR1-Scarlet*_diag_ext_F** | GATCAAAAGAGTGGTGAAATCAAA |
| ***PBR1-Scarlet*_diag_ext_R** | GGGCTCAAGGGTAACTCTTCTTT |

Capital letters represent genomic sequence

Small letters represent plasmid sequences, a linker, or a restriction site added

sgRNA_R: P2 (Min et al. 2016)

sgRNA_F: P3 (Min et al. 2016)

**Plasmids**

pV1093

CIP10-mScarlet-IDT

pFA-ARG4

pFA-GFP-CAURA3

pFA-GFP-HIS1
